# The Illusion of Polygenicity in Pool-seq Genetic Mapping studies: Insufficient Power Can Mask Simple Genetic Architectures

**DOI:** 10.1101/2025.07.23.666414

**Authors:** Anthony D. Long, Katherine M. Hanson, Stuart J Macdonald

## Abstract

Pool-seq (pooled sequencing) combines DNA from multiple individuals prior to sequencing, enabling population-level allele frequency estimation without individual genotyping. When employed in Genome Wide Association Studies (GWAS) pool-seq faces a fundamental power limitation in that errors on allele frequency estimates are proportional to sequence coverage. Although this power limitation is widely appreciated, pool-seq GWAS lacking unambiguous hits are often interpreted as showing a highly polygenic genetic architecture. We illustrate the limitation of inferring architecture from Manhattan plots using empirical data from a *Drosophila* zinc resistance mapping study. Despite achieving an average of >700× sequencing coverage in case and control pools, a directly ascertained SNP-based GWAS failed to reveal clear evidence for major-effect loci. A unique feature of the dataset is that an advanced intercross multiparent population, with known founders, was employed as the base population for the GWAS. We leverage this unique population structure to carry out a second GWAS using imputed haplotype frequency estimates, which in contrast revealed localized regions of major effect. A third reanalysis of the same data using imputed SNP genotypes derived from the founder haplotype frequency estimates uncovered a similar major gene architecture. The key difference between approaches lies in statistical power: directly ascertained SNP counts have errors proportional to sequencing coverage whereas known founder imputation-based approaches can be considerably more accurate. This work highlights that insufficiently powered GWAS studies can mask simple genetic architectures and create the illusion of polygenicity through statistical noise alone.

---

Pool-seq (pooled sequencing) is a cost-effective genomic approach where DNA from multiple individuals is combined prior to short-read sequencing, allowing researchers to estimate the allele frequency in a population without individually genotyping each individual (Futschik and Schlötterer 2010; Schlötterer et al. 2014). Pool-seq has been widely employed across diverse research contexts including population genomics (Kofler et al. 2011; Czech et al. 2024), experimental evolution studies (Burke et al. 2010; Turner et al. 2011; Orozco-terWengel et al. 2012; Burke et al. 2014), agricultural breeding programs (Michelmore et al. 1991), and genetic mapping in model organisms (Ehrenreich et al. 2010; Macdonald et al. 2022). A fundamental principle of pool-seq is that, for autosomal loci, when the number of chromosomes in the pool, 2*N*, substantially exceeds the depth of sequence coverage, *C* (*i*.*e*., *2N >> C*), directly ascertained SNP frequency estimates provide unbiased estimates of allele frequencies in the population from which the sample is drawn (Futschik and Schlötterer 2010). Furthermore, the precision of these frequency estimates follows binomial sampling theory, with standard errors proportional to *√(pq/C)*, where *p* and *q* are the reference and alternative allele frequencies.

Pool-seq has been increasingly applied to case-control genome-wide association studies (GWAS), where researchers compare allele frequencies between phenotypically distinct pools of individuals to identify genetic loci contributing to complex trait variation (Huang et al. 2012; Bastide et al. 2013; Morozova et al. 2015; Fochler et al. 2017; Zhou et al. 2017; Macdonald et al. 2022; Macdonald and Long 2022). This approach is analogous to the human case-control GWAS design (Risch and Merikangas 1996; Wellcome Trust Case Control Consortium 2007), in which a phenotypically extreme group of *N*_*1*_ affected case individuals are compared to *N*_*2*_ control individuals, except that rather than genotyping individuals via SNP arrays, pools of DNA from case or control individuals are subjected to short read sequencing to coverages *C*_*1*_ and *C*_*2*_. The standard GWAS tests for differences in allele frequencies between the *2N*_*1*_ case versus *2N*_*2*_ control autosomal alleles, whereas under the pool-seq design the tests are for differences between the *C*_*1*_ case versus *C*_*2*_ control reads per locus. Several studies have successfully implemented this approach across model organisms demonstrating its feasibility for trait dissection (Craig et al. 2009; Bastide et al. 2013; Lirakis et al. 2022; Davies and Myles 2023). However, much like a standard GWAS, pool-seq GWAS faces the fundamental trade-off between controlling false positive rates through stringent significance thresholds and maintaining sufficient statistical power to detect true associations. For example, The Wellcome Trust Case Control Consortium established that human GWAS requires significance thresholds of at least -log10(p-value) > 6, paired with on the order of three thousand fully genotyped case versus control individuals, to reliably detect subtle effect risk alleles while controlling for multiple tests (Wellcome Trust Case Control Consortium 2007).

The power limitations of case-control comparisons using pool-seq are well-established (Baldwin-Brown et al. 2014; Kofler and Schlötterer 2014): to reliably detect allele frequency differences <10%, pools of thousands of individuals sequenced to >1000X average coverage are likely required. Despite this constraint, Manhattan plots of -log10(p-value) against genomic location from experiments with much lower sequence coverage, and that show little evidence of “hits”, are often interpreted as representing a highly polygenic architecture (Huang et al. 2012; Orozco-terWengel et al. 2012; Morozova et al. 2015; Fochler et al. 2017; Zhou et al. 2017; Barghi et al. 2019; Lirakis et al. 2022). A polygenic architecture may indeed underlie the trait, but Manhattan plots from modestly powered studies are a poor proxy for genetic architecture. To quantify this limitation, we conducted simulations of pooled sequencing case-control experiments with pools of 10,000 individuals, varying levels of sequencing coverage (400X, 1000X and 5000X), and a single SNP with a true allele frequency difference of 4% or 8% between pools (Figure 1). Such allele frequency differences would be considered large by the standards of human disease GWAS studies, yet the sample sizes and coverages required to detect these loci are never achieved in published pool-seq studies. Our simulations suggest that only at 5000X coverage for an allele frequency difference of 8% is the allele of large effect reliably detected. It would be easy to conclude from the Manhattan plots for the more modestly powered experiments shown in Figure 1 that the simulated trait has a polygenic basis, and that many of the sites with highest LOD scores are true, but very modest-effect causative loci. The combination of limited power and genome-wide testing can generate apparent polygenic signals through statistical noise alone. This phenomenon, where insufficient statistical power masks a relatively simple genetic architecture, represents a critical interpretive challenge in pool-seq studies.

**Figure 1:**
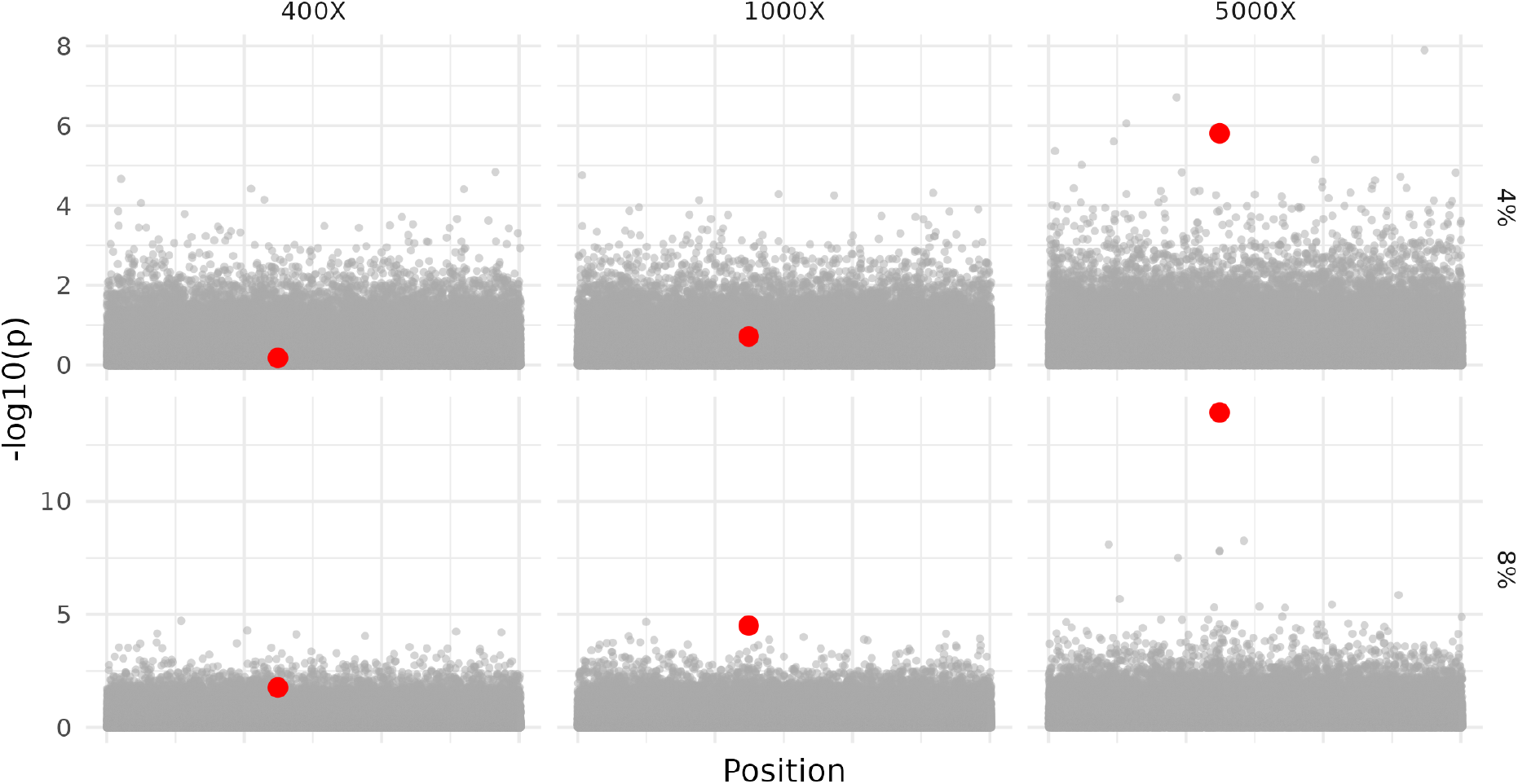
Simulated pool-seq case/control experiments. We simulated 500 haplotypes from a 1Mb region with typical *D. melanogaster* population genetic parameters. These haplotypes were instantaneously expanded to create 10,000 allele Case versus Control pools, subsequently pool-sequenced to expected coverages of 400X, 1000X, or 5000X. The case haplotype pools were sampled conditioning on a single intermediate frequency focal SNP (labelled in red) having a true allele frequency difference between cases or controls of 4% or 8%. We simulate a 1Mb region here and note a -log10(p) threshold of 5 is commonly used in *D. melanogaster* GWAS studies for statistical significance.

A recently published dataset in *Drosophila melanogaster* allows us to uniquely illustrate the problem in an empirical as opposed to a simulated dataset. Hanson et al. (Hanson et al. 2025) utilized a base population derived by combining 663 Drosophila Synthetic Population Resource (DSPR) 8-way, known founder advanced intercross recombinant inbred lines. They collected 0-24 hour old embryos and exposed them to either 25 mM zinc chloride or water. Case samples were generated from the ∼7% most zinc-resistant adult females that survived the zinc treatment, while controls represent samples of individuals that developed successfully in normal media conditions. Across 12 replicate selection experiments, a total of 4831 control and 3448 case individuals were sampled and pooled for sequencing. A total of 739× and 983× sequence coverage for the zinc-treated “case” and water-treated control pools was obtained, respectively (Figure 2). Ignoring the unique population structure of the base population employed, the experiment is a large case-control GWAS by Drosophila standards. Indeed, it is conceivable that the power of the experiment exceeds that of a typical 200-500 inbred-line-based mapping study (Macdonald et al. 2022).

**Figure 2:**
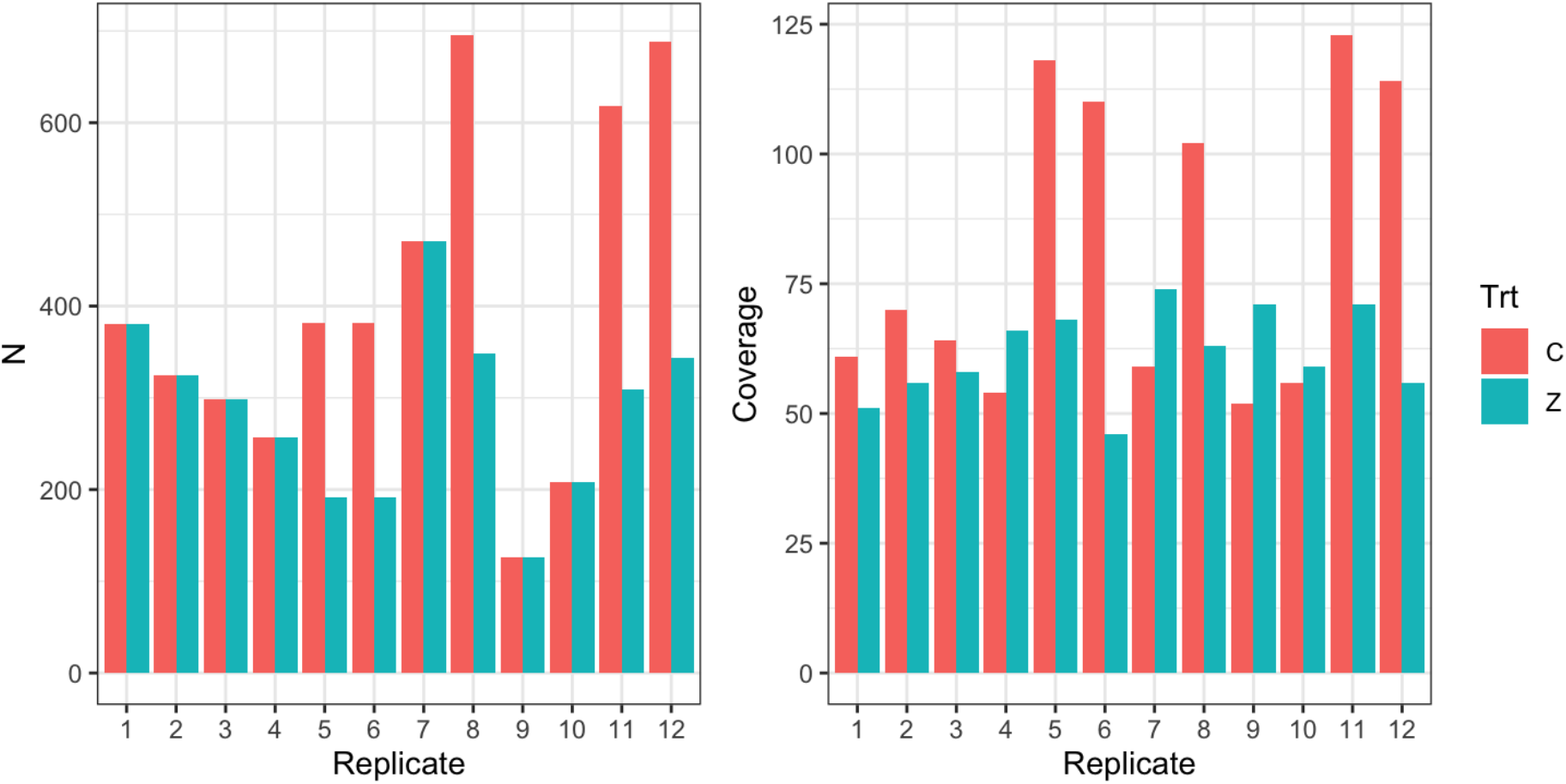
Sampling properties of an X-QTL experiment challenging a *Drosophila melanogaster* synthetic population with zinc chloride (Hanson et al. 2025). The experiment consisted of 12 paired replicates of <u>C</u>ontrol versus <u>Z</u>inc-selected treatments (Trt). For each treatment and replicate N individuals were sampled to create DNA pools (left panel), and these were subsequently pool-sequenced to the coverage depicted in the right panel. Replicates 5, 6, 8, 11 and 12 had two separate, equally sized control pools.

We carried out a GWAS of these data by contrasting directly ascertained allele frequencies between cases and controls, summarizing the experiment with a Manhattan plot (Figure 3A). Although 642 SNPs exceed a Bonferroni significance threshold, these significant associations show little apparent localization to specific genomic regions and instead appear dispersed throughout the genome. Based on these data alone one could easily conclude that zinc resistance in this population has a highly polygenic genetic architecture.

**Figure 3:**
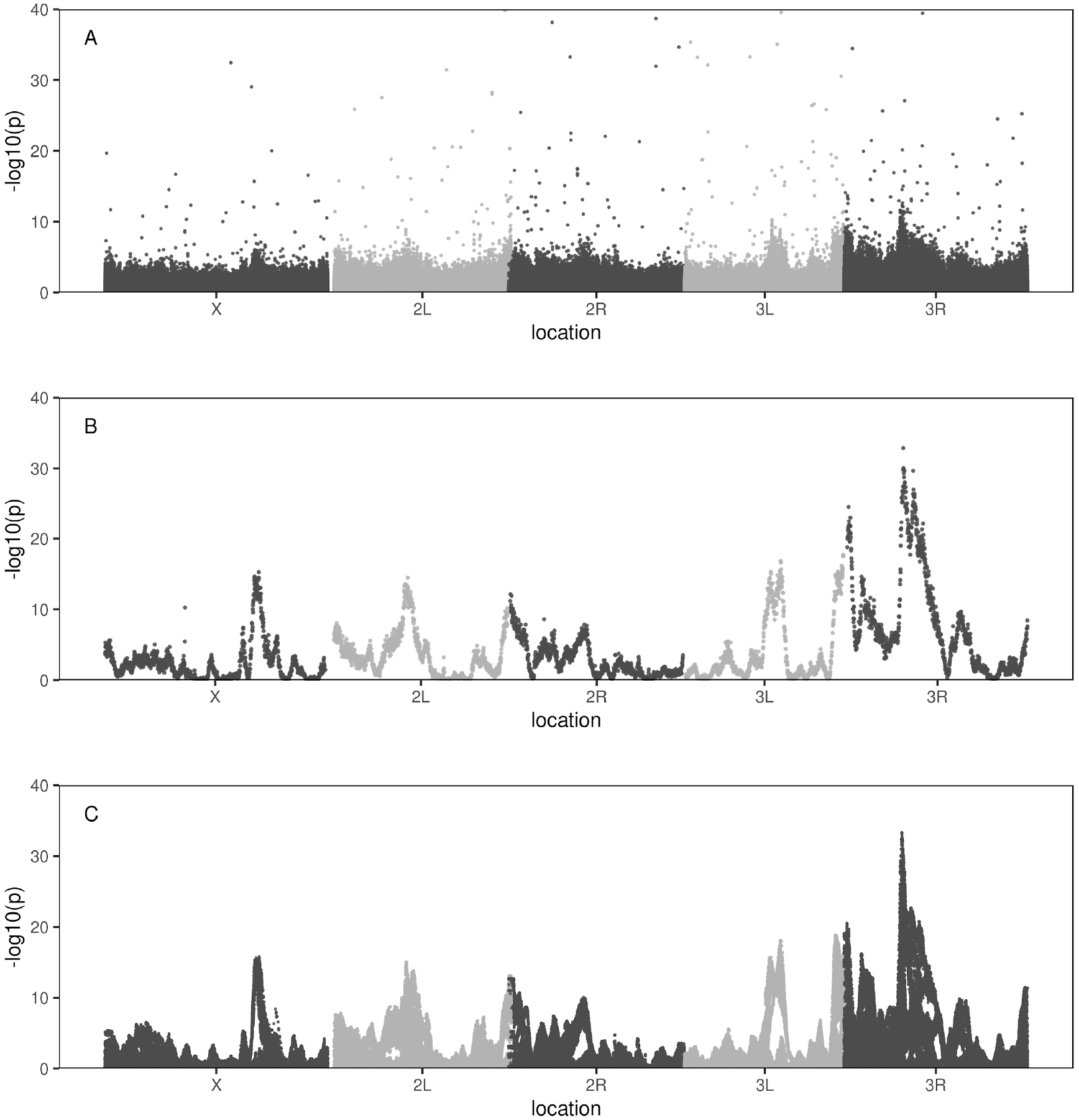
Zinc chloride developmental resistance Manhattan plots associated with three different approaches to statistical testing: (A) directly ascertained REF and ALT SNP counts (we do not display a statistical threshold but note a -log10(p) threshold of 5 is widely used in Drosophila GWAS), (B) Imputed founder haplotype counts, and (C) imputed SNP counts. X-axes are scaled in genetic as opposed to physical distances. *p*-values are obtained from Cochran–Mantel–Haenszel tests on counts over the 12 replicates of the experiment. For directly ascertained SNPs, the counts used are observed REF vs ALT counts. For the imputation-based tests, *pseudo*-counts are the product of haplotype (or SNP) frequency estimates and twice the number of individuals per DNA pool. These pseudo-counts will inflate -log10(p-values) by a constant multiplicative factor.

The dataset of Hanson et al. (Hanson et al. 2025) is unique in that the base population employed is derived from an advanced intercross synthetic population originating from eight known founder strains. This experimental design allows the frequency of those eight founder alleles to be estimated from the pool-seq data throughout the genome using a sliding window haplotype estimator (Long et al. 2011; Kessner et al. 2013; Burke et al. 2014; Tilk et al. 2019; Linder et al. 2020; Linder et al. 2022; Macdonald et al. 2022). Specifically, Hanson et al. estimated founder haplotype frequencies in overlapping 1.5-cM windows across the genome for each sample. These eight founder haplotype frequencies were then tested for differentiation as a function of genomic location. This known-founder haplotype-based testing strategy underlies the X-QTL (or bulked segregant) analysis approach to mapping QTL (Ehrenreich et al. 2012; Macdonald et al. 2022; Macdonald and Long 2022). Based on this X-QTL statistical approach, employing the same raw data as the directly ascertained SNP GWAS described above, Hanson et al. concluded that resistance is associated with regions of major effect localized at centimorgan scales. They further identified and validated several major genes in these intervals (including *MTF-1*, a gene central to zinc homeostasis). Figure 3B reproduces the scan of Hanson et al and clearly shows highly significant peaks of association localized to specific genomic regions. The results of this zinc-resistance mapping are consistent with other studies where multiparent population (MPP)-based mapping have identified regions of large effect (King et al. 2012; King et al. 2014; Najarro et al. 2015; Everman et al. 2021). They present a stark contrast to the dispersed association pattern observed with directly ascertained SNP-based analysis.

We claim that the fundamental reason the haplotype-based test succeeded in detecting major-effect loci, while the direct SNP analysis failed, is simply due to poor power of the latter approach. Despite achieving greater than 700× sequencing coverage per treatment, the directly ascertained SNP approach still suffers from relatively large binomial sampling errors on the SNP frequency estimates. In turn, this noise in allele frequency estimation obscures genuine allele frequency differences between pools. In contrast, the haplotype-based approach leverages the critical statistical advantage that haplotype frequency estimates have much smaller errors than directly ascertained SNP frequencies. The error on imputed haplotype frequencies are proportional to ∼75-100% of the binomial sampling errors on twice the number of individuals per treatment, as opposed to the average short-read coverage (Tilk et al. 2019; Linder et al. 2020). In the zinc chloride experiment the average number of autosomal alleles per treatment (2×4140 = 8280) is roughly ten-fold greater than the average sequence coverage per treatment (700×), and as a result, haplotype frequencies are estimated with much greater certainty than directly ascertained SNP frequencies. The haplotype approach effectively circumvents the coverage limitation of directly ascertaining SNP frequencies. At sequencing coverages much less than 2N, as is typically the case with pool-seq datasets, imputing founder haplotype frequencies is a “trick” that can be used to achieve frequency estimation errors approaching genotyping every individual.

To illustrate that the difference we observe between the haplotype-based approach (Figure 3B) and the directly ascertained SNP-based approach (Figure 3A) is primarily due to sampling error on frequency estimates, and not somehow associated with the decision to employ haplotypes or some other detail of X-QTL mapping procedure, we used the haplotype frequency estimates to impute SNP frequencies. That is, we imputed SNP frequencies throughout the genome as the sum of the Hadamard product of the founder frequency estimates and known SNP states in the founders at any given location. Although counter-intuitive, just like the haplotype frequencies, these imputed SNP frequencies are estimated much more accurately than the directly ascertained SNP counts (Tilk et al. 2019; Linder et al. 2020). We then converted imputed SNP frequencies to pseudo-counts based on the number of individuals in each pool and carried out a third GWAS on the imputed SNP counts (Figure 3C). The plot demonstrates that when we use the more accurate imputed, as opposed to directly ascertained, SNP allele frequencies we recover the clear peaks of association seen in the haplotype-based analysis. This confirms that the failure of direct SNP analysis stems from poor power. Looking closely at the tests on imputed SNP frequencies (Supplementary Figure 1) it further appears as though one could fit several lines to that data. This pattern represents sets of SNPs that are “in phase” with one another with respect to different founder chromosomes and is expected in multiparent panels derived from a modest number of founders.

The power limitations we demonstrate are not specific to synthetic populations or X-QTL methodology but represent a fundamental challenge for pooled sequencing approaches. While our analysis employed a synthetic population derived from eight known founders, the underlying statistical problem, that errors on directly ascertained SNP frequency estimates are large relative to subtle allele frequency differences between pools, applies broadly to any pooled sequencing design. In outbred populations lacking known founder structure, researchers cannot employ the imputation strategy that rescued power in the illustrative example we provide here. The apparent success of haplotype-based approaches in synthetic populations stems not from any inherent superiority of haplotypes over SNPs, but simply because haplotype frequencies are estimated more accurately, effectively circumventing the coverage limitations that plague direct SNP ascertainment.

A criticism that can be leveled at the illustrative example presented in this work is that the statistical tests for directly ascertained SNPs use total counts derived from sequence coverage (see Methods), whereas the two imputation-based tests use total counts associated with twice the number of individuals in the pool from which DNA is made. Counts associated with the 2N individuals in the DNA pools result in higher power, since 2N is more than ten times the sequence coverage per pool. However, the total sequence coverage is the true count associated with directly ascertained SNPs, whereas the 2N associated with imputation-based tests incorrectly assumes imputation is equivalent to perfectly genotyping the N individuals in each pool. Imputation is simply not that effective. But the decision to multiply frequencies by 2N to obtain counts *only* impacts the statistical test as a multiplicative constant throughout the genome. As a result, the imputation-based Manhattan plots of Figure 3B&C simply have an “inflated” Y-axis scale. But this inflation constant does not contribute to the noise of the directly ascertained GWAS.

A second criticism is that an MPP is not appropriate for a GWAS-type analysis, since a synthetic population derived from an 8-way cross is not the type of population typically employed in a GWAS study (generally a sample of natural chromosomes lacking a known set of founders). This is reflected in Figure 3 where for the two imputation-based scans the “hits” extend over centimorgan sized intervals, a much larger region than the localization signal expected of a GWAS. But this difference doesn’t detract from the value of our illustration. We observe that directly ascertained SNP frequencies at coverages approaching 700X per sample are insufficient to detect the real, but subtle frequency shifts apparent in this dataset, a result independent of the genetic details of the population employed. If anything, the problem is more dire in a population harboring lower levels of linkage disequilibrium as a function of distance. In such a case one is trying to detect the signal associated with only a handful of SNPs in LD with a causative factor, and a lower genome-wide *p*-value threshold is likely required to control for false positives.

In *Drosophila melanogaster*, we are aware of four traits have been dissected using both an MPP-panel based mapping approach using the *Drosophila* Synthetic Population Resource (DSPR; King et al. 2012) and a GWAS-based analysis using collection of inbred lines called the *Drosophila* Genetic Reference panel (DGRP; Mackay et al. 2012; Huang et al. 2014) where the genetic architecture appears complex: resistance to caffeine (Najarro et al. 2015), copper toxicity (Everman et al. 2021; Everman et al. 2023), boric acid resistance (Najarro et al. 2017), and starvation stress (Everman et al. 2019). In all four cases DSPR mapping-based approaches have successfully identified loci of large effect, while in contrast DGRP association-based approaches have concluded that trait variation is highly polygenic and due to dozens to hundreds of loci of subtle effect spread throughout the genome. Notably the association hits are subtle enough that they rely on a suggestive threshold for significance, often a -log10(p-value) > 5. Our observations with respect to pool-seq case-control designs suggest that power deficits can lead to incorrect assumptions about the polygenicity of complex traits, and we speculate that a similar pattern may explain the contradictory results observed in these inbred line studies. The highly polygenic signal in these four association studies is consistent with DGRP-based GWAS outcomes more generally (Mackay and Huang 2018), as well as those in other Drosophila GWAS-type studies employing individual-based genotyping (Pallares et al. 2023). Perhaps the observed difference between DGRP and DSPR outcomes is similarly driven by differences in power. Overall, in Drosophila the genetic basis of trait variation dissected using GWAS-type approaches appears highly polygenic, while in contrast traits dissected using QTL-mapping type approaches appear more oligogenic, and this remains true even when holding the trait constant. It is possible that the apparent contradiction between polygenicity versus handfuls of more major effect loci is driven by power. Many traits currently characterized as highly polygenic based on dispersed GWAS signals are consistent with the traits truly being polygenic, but equally consistent with a more oligogenic architecture that studies simply are poorly powered to detect.

## Methods

The zinc chloride mapping dataset is described in detail in Hanson et al. (Hanson et al. 2025). Briefly, the study utilized a population derived by intercrossing 663 Drosophila Synthetic Population Resource (DSPR) “A” founder population recombinant inbred lines. After 31-36 generations of synthetic population maintenance, 12 replicate zinc selection assays were performed. To perform the assays, 0–24-hour old embryos were collected from the base population and transferred to control (water) or treatment (25-mM zinc chloride) bottles. Emerged adult females were collected, and pooled DNA samples were prepared for both control and zinc-selected groups, DNA libraries were prepared using the Illumina DNA Prep protocol and sequenced with PE150 reads on an Illumina NovaSeq 6000. Reads were mapped to the *D. melanogaster* Release 6 reference genome, REF/ALT counts were obtained for every SNP, and founder haplotype frequencies estimated in overlapping 1.5-cM windows across the genome for each sample.

Here, we employ the raw SNP tables and haplotype frequency estimates from Hanson et al. (2025). But unlike that earlier work all the statistical tests in the present study employ Cochran– Mantel–Haenszel (CHM) tests on counts as opposed to ANOVA on square root transformed frequency estimates. The zinc chloride experiment consisted of twelve replicate control versus zinc-selected comparisons, so CHM is an appropriate test. Additionally, the use of the CHM test for all three analytical approaches allows for more intuitive comparisons between the study designs. For directly ascertained SNP scans the CHM tests employed raw REF vs ALT counts tables, and as a result the total counts in the analysis at any given SNP is the total coverage at the SNP. In contrast the two imputation-based designs use the raw counts and founder genotypes to impute haplotype or SNP frequencies at 10kb steps, or at each SNP in the genome, respectively. SNP frequencies are imputed by multiplying the closest set of haplotype frequency estimates by the founder states (generally 0 or 1) for each haplotype. To convert per-sample imputed haplotype or SNP frequencies to counts, those frequencies are multiplied by twice the number of individuals contributing to that pooled DNA sample. The number of individuals in a pool is a genome-wide constant per pool, as a result it will not generate variation between sites, instead it will result in -log10(p-values)’s inflated by a constant factor.

MSPrime (Kelleher and Lohse 2020; Baumdicker et al. 2022) was used to simulate 500 haplotypes from a neutrally evolving 1Mb region under *Drosophila melanogaster*-like parameters (*N*_*e*_=1e6, *u*=5e-9, *r*=2e-8). These haplotypes were sampled with replacement to create a ten thousand haplotype control base population. A similar 10,000 haplotype case population was created by sampling haplotypes with replacement from the control population conditional on a single focal SNP having an allele frequency difference of 4% or 8% from the controls. Frequencies were then estimated at all common SNPs for both case and control populations, and then pool-seq allele counts obtained via random negative binomial draws (size = mu = expected coverage) conditional on the known frequencies and expected sequencing coverages of 400X, 1000X, or 5000X. The negative binomial was employed so that realized coverage varied among SNP positions according to an over-dispersed poisson with a variance twice the mean (equal to expected coverage). Chi-square tests were carried out at each SNP and -log10(p-values) obtained.

## Data and Methods Availability

The raw data, methods for calling SNPs, and haplotyping calling are described in the Hanson 2025. Briefly, DSPR RILs are available from the BDSC, raw FASTQ files are in the NCBI SRA under accession number PRJNA1127662, and all code to replicate the analyses is available on GitHub (https://github.com/Hanson19/Zinc-X-QTL). The analysis code, and code to reproduce the figures of this work are hosted on Github (https://github.com/tdlong/Compare_3_GWAS.git).

## Acknowledgements

This work was supported by NIH award R01-OD034064 (to SJM and ADL).

---

**Supplementary Figure 1:**
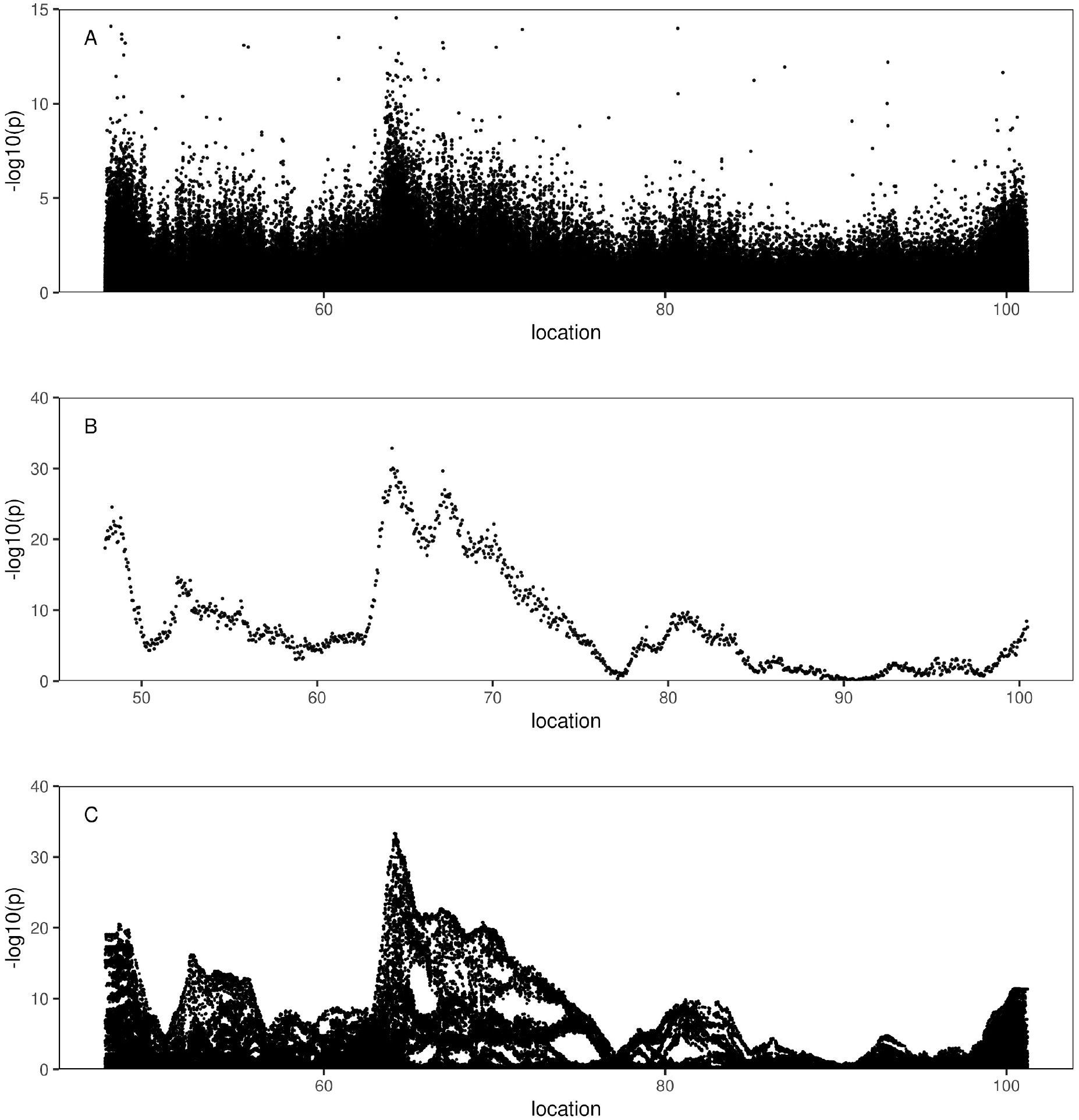
Zinc chloride developmental resistance Manhattan plots associated with three different approaches to statistical testing for chromosome 3R: (A) directly ascertained REF and ALT SNP counts, (B) Imputed founder haplotype counts, and (C) imputed SNP counts. These panels are otherwise the same as Figure 3, except for panel A where we limit the Y-axis to 15.

